# Where is ADHD in the brain? Evidence for a neurodevelopmental continuum of brain dynamics

**DOI:** 10.64898/2026.05.04.722738

**Authors:** Gian Marco Duma, Giulia Stefanelli, Lisa Toffoli, Giulia Ferri, Giovanni Pellegrino, Alberto Danieli, Federica Martinez, Vincenza Tarantino, Duncan Astle, Fiorella Del Popolo Cristaldi, Giovanni Mento

**Affiliations:** Scientific Institute IRCCS E. Medea, Conegliano (TV), Italy; NeuroDev Lab, Department of General Psychology, University of Padova, Padova, Italy; Department of Electronic System, Aalborg University, Denmark; Department of Clinical Neurological Sciences, Schulich School of Medicine and Dentistry, Western University, London, ON, Canada; Neuropsychology Lab, Department of Psychology, Educational Sciences and Human Movement, University of Palermo, Palermo, Italy; MRC Cognition and Brain Sciences Unit, University of Cambridge, Cambridge, United Kingdom; Department of Psychiatry, University of Cambridge, Cambridge, United Kingdom

**Keywords:** ADHD, Brain dynamics, Cortical excitability, Brain–behaviour relationships, Neurodevelopmental dimensions

## Abstract

**Background:** Attention-deficit/hyperactivity disorder (ADHD) has traditionally been conceptualized categorically, with efforts to identify disorder-specific neurobiological endophenotypes. However, dimensional models suggest that brain–behavior organization may follow developmental axes that cut across diagnostic boundaries. We tested whether neural dynamics and cortical excitability differentiate those with ADHD diagnoses from typically developing (TD) peers, and whether brain–behavior covariance aligns with diagnostic or developmental dimensions.

**Methods:** We studied 84 participants aged 8–17 years (51 ADHD, 33 TD). High-density electrophysiological (hdEEG) measures included task-free source-resolved data used to derive mean global brain fluidity (variance of dynamic functional connectivity) and region-specific cortical excitability. Behavioural measures included self- and parent-report questionnaires, cognitive control (CC) tasks, and neuropsychological tests. Partial least squares (PLS) assessed multivariate brain–behavior associations including age, followed by clustering based on latent component scores.

**Results:** Group differences emerged in parent-report questionnaires and CC tasks, but not in neuropsychological measures. ADHD individuals showed higher mean global brain fluidity and increased cortical excitability. The excitability–fluidity relationship was network-dependent: higher excitability predicted higher fluidity in task-positive networks and lower fluidity in default-mode and salience networks, with no group effects. PLS identified a latent dimension linking neural metrics with age, verbal fluency, inhibitory control, and positive affect, but it did not distinguish ADHD from TD. Clustering revealed two neurodevelopmental profiles spanning both groups.

**Conclusions:** While ADHD is associated with mean-level differences in neural dynamics, brain–behaviour organization follows a developmental neurocognitive–affective axis that transcends the diagnostic boundary. These findings support a dimensional framework for understanding neurobiological variation in neurodevelopmental conditions.

---

Attention-Deficit/Hyperactivity Disorder (ADHD) is a neurodevelopmental condition with a global prevalence of 4–8% in children and adolescents (Cortese et al., 2023; Jackson et al., 2025), typically emerging in middle childhood and often persisting into later developmental stages (Franke et al., 2018; Posner et al., 2020). It is characterized by developmentally inappropriate levels of inattention and/or hyperactivity-impulsivity that interfere with academic, social, and emotional functioning (Faraone et al., 2024; Sonuga-Barke et al., 2023). Core behavioural features include difficulties in sustained attention, response inhibition, motor regulation, and goal-directed organisation, mapping onto the domains of inattention, hyperactivity, and impulsivity(Arabacı & Parris, 2020; Barkley et al., 2002; Cortese, 2020; Faraone et al., 2024). These manifestations are unlikely to be attributable a singular underlying cause but reflect broader alterations in cognitive control (CC) and self-regulation processes (Faraone et al., 2015; Nigg et al., 2018; Posner et al., 2020; E. J. S. Sonuga-Barke & Castellanos, 2007). Within this framework, executive functions (EFs), such as working memory, cognitive flexibility, inhibitory control, and planning, are commonly implicated and have been linked to altered fronto-striatal and cerebellar circuitry (Barkley, 1997; Castellanos & Tannock, 2002; Nigg & Casey, 2005; Pennington & Ozonoff, 1996; Schachar et al., 2000). However, EF difficulties alone do not fully explain all characteristics of those with the condition (Willcutt et al., 2005). Altered reward processing, motivation, emotional reactivity, and self-regulation have been consistently reported in ADHD (Faraone & Larsson, 2019; E. Sonuga-Barke et al., 2010). While these processes are sometimes subsumed under broader conceptualisations of executive functioning, they are also frequently examined as partially distinct components of self-regulatory functioning, supporting a heterogeneous and dimensional view of ADHD (Astle et al., 2022; Sonuga-Barke et al., 2023).

Higher-order cognitive processes, particularly CC, are supported by the coordinated organization of large-scale brain networks, including the central executive network (centred on dorsolateral prefrontal and posterior parietal cortices), the salience network (mainly anchored in anterior insula and dorsal anterior cingulate cortex), and the default mode network (comprising medial prefrontal, posterior cingulate/precuneus, and lateral temporoparietal regions) (Bassett & Sporns, 2017; Cohen & D’Esposito, 2016). Optimal network functioning has been linked to the balance between excitatory and inhibitory neural signaling, which is thought to constrain how neural systems transition between functional states, thereby influencing both network integration and segregation. In ADHD, alterations have been observed both within and between these networks, particularly those supporting higher-order cognitive functions such as CC (Michael et al., 2025; Yin et al., 2022). These findings suggest that disrupted coordination across large-scale networks may contribute to the characteristic impairments in cognitive control and self-regulation.

At a more mechanistic level, optimal brain function has been linked to the balance between excitatory and inhibitory (E/I) neural signaling, which constrains how neural systems transition between functional states and shapes both network integration and segregation. Variability in neural excitability may therefore provide a bridge between local neural dynamics and large-scale network organization, with downstream implications for individual differences in CC and self-regulation (Gao et al., 2025). Disruptions in E/I balance, particularly involving GABA-mediated inhibition, have been broadly implicated in neurodevelopmental conditions and may contribute to the heterogeneous behavioural profiles observed in ADHD, where reduced GABA levels and altered inhibitory signaling have been reported (Puts et al., 2020; Richter et al., 2007; Sohal & Rubenstein, 2019; Sylvester et al., 2025). Such alterations may contribute not only to atypical network organization but also to behavioral dysregulation.

Despite the identification of numerous neurophysiological differences, including variations in oscillatory activity, connectivity patterns, and E/I balance indices, no single biomarker has demonstrated sufficient specificity or sensitivity to serve as a reliable diagnostic indicator of ADHD (Chen et al., 2024; Cortese et al., 2021, 2023; Li et al., 2021). Attempts to define a stable “brain fingerprint” uniquely characterizing ADHD have been limited by substantial inter-individual variability and by overlapping neural features with typically developing (TD) populations and other neurodevelopmental disorders. Indeed, similar alterations in connectivity or oscillatory dynamics can be observed across different clinical conditions, challenging the notion of a disorder-specific neural signature (Astle et al., 2022; Sonuga-Barke et al., 2023). These considerations support the growing push towards a transdiagnostic framework for understanding neurodevelopmental conditions. Rather than focusing on categorical diagnostic labels, this transdiagnostic framework seeks to identify dimensional functional profiles that cut across traditional boundaries (Astle et al., 2019; Fletcher-Watson, 2022; Siugzdaite et al., 2020; Sonuga-Barke et al., 2016, 2023). Put simply, given the marked variability within ADHD and the substantial overlap with other neurodevelopmental conditions, as well as with TD individuals, it is unlikely that a single neural signature defines this diagnostic group (Cortese et al., 2021; Siugzdaite et al., 2020). Instead, integrating neurofunctional patterns with specific cognitive and behavioural dimensions may offer a more comprehensive and reliable understanding of neurodevelopmental conditions and provide a clinically informative framework. In the present study, we aimed to characterize the neurofunctional profile of ADHD by integrating task-free neural measures with behavioural and cognitive dimensions targeting core symptoms. To capture intrinsic large-scale brain dynamics, we focused on two electroenchephalography (EEG)-derived indices, applied here for the first time in ADHD: (1) an excitability index (EI), capturing local excitability level from intrinsic brain activity, and (2) a fluidity index (FI), characterizing the temporal variability of large scale brain dynamics. Furthermore, we also examined their interaction to assess how local excitability relates to the stability of large-scale dynamics in individuals with ADHD.

Participants with ADHD and a typically developing (TD) comparison group underwent comprehensive characterization of behavioural, cognitive, and emotional domains, using standardized neuropsychological tests alongside self- and parent-report questionnaires assessing behavioural patterns, emotional regulation, and everyday functioning. By combining multi-level behavioural data with cutting-edge, EEG-derived metrics, we sought to delineate an integrated neurofunctional profile. Our overarching aim was to characterize how variability in neural excitability and large-scale brain dynamics relates to individual differences in cognitive and behavioural functioning in ADHD. Specifically, we pursued three objectives: (i) to test whether these neurofunctional signatures distinguish individuals with ADHD from typically developing controls; (ii) to examine whether they track key symptom dimensions; and (iii) to determine whether these associations are specific to diagnostic status or reflect broader neurodevelopmental variation.

## Method

### Participants

We initially enrolled 99 children and adolescents (26 females) aged between 8 and 17 years. Participants were recruited from primary and secondary schools in the Veneto region (Italy), as well as from a database managed by the NeuroDev laboratory of the University of Padova. This database was developed through collaborations with specialized clinical centers and family associations. For all participants, high-density EEG data (hdEEG) were collected at the Department of General Psychology (University of Padova), or at the Epilepsy and Clinical Neurophysiology Unit, IRCCS Eugenio Medea (Conegliano, Italy). In addition, participants underwent a neuropsychological assessment including an experimental task (Gonthier et al., 2021; Toffoli et al., 2025), whilst parents completed standardized behavioural questionnaires. Exclusion criteria and details of excluded participants (n=15) are reported in the Supporting Information. The final sample consisted of 84 participants, including 51 ADHD and 33 TD. Further details on the final sample are reported in Table 1. All individuals with an ADHD diagnosis received that diagnosis from a child psychiatrist and/and/or psychologist.

**Table 1.** Reports of the demographic characteristics of the sample across groups, including age mean and standard deviation (in months and years), sex distribution and age range. No difference in age (months and years) were identified between the two groups (all *p* > .05).

| Group | Sex |  | Mean ± SD | Mean ± SD | Range |
| --- | --- | --- | --- | --- | --- |
|  | Female | Male | (age in months) | (age in years) | (age in months) |
| Control | 15 | 18 | 140.58 ± 29.08 | 11.30 ± 2.42 | 97-204 |
| ADHD | 8 | 43 | 152.75 ± 27.77 | 12.25 ± 2.28 | 102-202 |

### Behavioural and Neuropsychological Assessment

Parents completed the questionnaires via an online platform (Qualtrics, 2019). Parent-report measures included the Conners’ Parent Rating Scale – Revised (CPRS-R; (Conners, 2001; Nobile et al., 2007), assessing behavioural and emotional difficulties; the Behavior Rating Inventory of Executive Function – Second Edition (BRIEF-2; (Gioia et al., 2000; Marano et al., 2016), measuring EFs in everyday contexts; and the Social Responsiveness Scale (SRS-2; (Constantino, 2013; D’Ardia et al., 2021), assessing social functioning.

Neuropsychological assessment was conducted in person and included working memory and strategic vocabulary access through backward digit span and phonemic fluency (Bisiacchi et al., 2005; Gugliotta, 2009). Cognitive flexibility and inhibitory control were assessed using the Wisconsin Card Sorting Test (WCST-64; (Heaton & PAR Staff., 2000)0), a modified version of the Flanker task (Gonthier et al., 2021; Toffoli et al., 2025), and the Inhibition subtest of the NEPSY-II (Urgesi et al., 2011). Participants also completed self-report questionnaires, including the Positive and Negative Affect Schedule – Short Form (PANAS-C; (Ciucci et al., 2017; Ebesutani et al., 2012; Thompson, 2007). Further details on the neuropsychological tests and questionnaires are provided in the Supporting Information.

### Task-Free EEG Recording

Seven minutes of task-free hdEEG was recorded with a R-Net EEG cap (128 electrodes; Brain Products®). Participants were seated comfortably on a chair in a silent room, looking at the Inscapes video (Vanderwal et al., 2015), which was specifically designed to improve compliance in developmental populations while minimizing motion and cognitive load during neural data acquisition (Vandewouw et al., 2021).

### EEG preprocessing

Signal preprocessing was performed using EEGLAB v2024.2 (Delorme & Makeig, 2004) following a pipeline validated in prior studies (Duma et al., 2022, 2023). First, data were downsampled to 250 Hz, bandpass-filtered (1-45 Hz), and then epoched (1 sec). Automated bad-channel and artifact detection and removal were performed using the Trial-by-Trial EEGLAB plugin, with the following parameters: channels differential average amplitude 150 μV in more than 30% of epochs. Epochs were removed if they contained more than 10 bad channels. We then used Independent Component Analysis to correct artifacts. We reduced the signal to 40 components and removed those clearly related by visual inspection to eye, muscle, channel, and heart artefacts.

### Cortical Source Modelling

Public, age-specific anatomical template images (Fonov et al., 2011) were segmented using FreeSurfer (Fischl, 2012) within the CBRAIN cluster (Sherif et al., 2014) and then imported into Brainstorm (Tadel et al., 2011). EEG electrodes were co-registered with the templates using standard anatomical landmarks, and manual adjustments were applied when necessary. The freesurfer-generated cortical surface mesh was downsampled to 15,002 vertices. The boundary element model (BEM) forward modelling was performed using OpenMEEG BEM (Gramfort, 2013; Kybic et al., 2005) and sLORETA (Pascual-Marqui, 2002) for the inverse solution (Brainstorm’s default parameter settings). Vertex-wise source activity was downsampled according to the Desikan-Killiany surface atlas (Desikan et al., 2006).

### Fluiditiy Index

Fluidity is a global measure of temporal stability of dynamic functional connectivity (FC), here extracted at the source space level. To calculate this, we first computed dynamic FC using lagged coherence (lagCoh). Unlike magnitude-squared coherence, lagCoh removes the instantaneous (zero-phase) component, thereby reducing the effects of volume conduction providing a more robust estimate of genuine time-delayed interactions (Chiarion et al., 2023; Hindriks, 2021; Nagy et al., 2024). Connectivity was computed with Brainstorm using a sliding-window approach (6 s window, 90% overlap) in the five canonical frequency bands: delta (2–4 Hz), theta (4–7 Hz), alpha (8–12 Hz), beta (13–30 Hz), and gamma (30–40 Hz), yielding a connectivity tensor of size N (region) × N × M (windows) × F (frequency). Successively, a dynamic functional connectivity matrix (dFC) was obtained, for each frequency, by correlating the upper triangular elements of lagCoh matrices from consecutive windows, capturing the temporal evolution of inter-regional synchronization (see Formulas 1 and 2 in Supporting Information). To quantify temporal variability in functional connectivity, we computed fluidity, a metric derived from the dFC matrices. Fluidity was defined as the variance of the upper triangular values of the dFC matrix after removing overlapping windows (see Formulas 1 to 5 in Supporting Information).

This metric reflects the temporal variability of functional connectivity and was interpreted as an index of the brain’s dynamic adaptability (Breyton et al., 2025).

### Excitability Index

The EI quantifies mean spatial phase synchronization and, particularly within gamma frequency bands, shows a strong correlation with cortical excitability as assessed using perturbational methods and as modulated by antiseizure medications in patients with epilepsy (Meisel et al., 2015, 2016). This excitation–inhibition measure is conceptually related to the widely used phase-locking value (PLV), with the key distinction that it was computed in the gamma band (30-40 Hz) across spatial dimensions rather than over time.

### Statistics

A first analysis was conducted using independent-samples t-tests to compare groups across all behavioural variables. For each variable, group means (± SD) and t-values are reported in the Table S2. False discovery rate (FDR) correction was applied to control for multiple comparisons across all variables(Benjamini & Hochberg, 1995).

Group differences in fluidity were assessed using a generalized linear model (GLM) in R ((R Core Team, 2020); version 4.4.0) with a Gamma distribution and log link function: Fluidity ∼ Group × Frequency bands + Age. Similarly, to identify local differences in excitability between groups, we fitted: Excitability ∼ Group × Brain_Region + Age. Subsequently, we examined whether excitability modulated fluidity differently between groups. To this end, for each cortical region and frequency band, we fitted a FDR-corrected mass-univariate linear model: log(Fluidity) ∼ Group × Excitability, with fluidity log-transformed to reduce skewness.

To investigate multivariate associations between behavioural measures and brain activity we performed a Partial Least Square (PLS) analysis (Krishnan et al., 2011) using MATLAB’s plsregress function (Rosipal & Krämer, 2006). This method identifies latent components that maximize shared covariance between two high-dimensional datasets, making it particularly suitable for examining distributed brain–behavior relationships in the presence of collinearity. Behavioural data (subjects × [behavioural variables + neuropsychological variables + age]) and brain data (subjects × [68 regional excitability values + 5 band-specific fluidity values]) were organised into separate matrices. Behavioral variables were z-scored across participants. The optimal number of latent variables (LVs) in the PLS model was determined using cross-validation to balance predictive performance and model complexity. Predictive accuracy was first assessed using leave-one-out cross-validation, where prediction error was quantified as cross-validated root mean squared error, and further validated using repeated 10-fold cross-validation (K = 10, R = 50), providing a more stable estimate of model generalizability (Kim, 2009; Wold et al., 2001). Both approaches consistently indicated a minimum prediction error for two components, with increasing error for higher model orders, suggesting a predominantly two-dimensional predictive structure and the onset of overfitting beyond the second component. Statistical significance was evaluated independently using permutation testing (10,000 permutations). The total variance explained in the brain matrix (R²Y) was used as a global index of model performance. At each iteration, the correspondence between behavioural and brain data was disrupted by randomly permuting subjects in the brain matrix, and the PLS model was recomputed. The observed total explained variance and component-specific effects were then compared against their respective null distributions. Component strength was additionally quantified as the correlation between behavioural and brain scores, with significance evaluated using a two-tailed permutation-based approach. To assess the reliability of individual behavioural and regional contributions, permutation-based inference was also applied to the loadings. For each permutation, component signs were aligned with the original solution to address sign indeterminacy, and empirical p-values were computed by comparing observed loadings to their null distributions. The analytical pipeline is represented in Fig.1.

**Figure 1.**
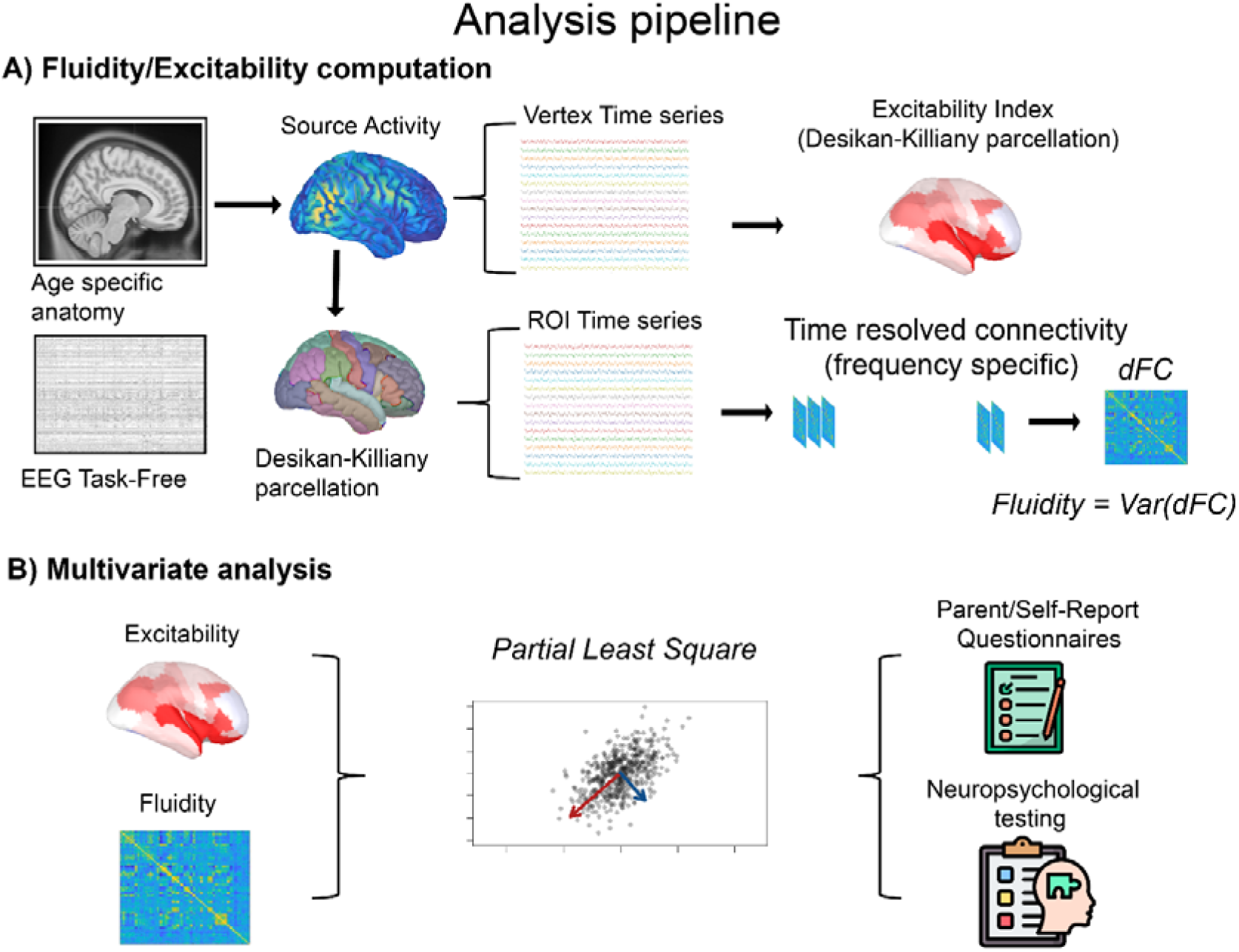
Analysis pipeline. Panel A: Overview of the analytical workflow, from age-specific magnetic resonance imaging templates and task-free EEG to source reconstruction. Subsequent steps include computation of the excitability index (defined as spatial phase coherence across vertices within each Desikan–Killiany atlas region), estimation of dynamic functional connectivity from atlas-downsampled source activity, and derivation of the fluidity metric. Panel B: Schematic representation of the neurofunctional and behavioral/neuropsychological measures entered into the brain–behavior analysis, performed using a multivariate approach based on partial least squares (PLS)

## Results

### Behavioural and Neuropsychological Assessment

Significant group differences were observed across all parent-report questionnaires, reflecting distinct patterns in behavioral regulation, executive functioning, and social responsiveness, with the ADHD group showing higher (i.e., worse) scores. Neuropsychological performance partly supported these findings. The ADHD group showed a reduced percentage of correct responses and conceptual level responses on the WCST-64, indicating reduced cognitive flexibility. Likewise, in the adaptive Flanker task, individuals with ADHD showed less efficient CC adaptation between high- and low-conflict blocks, as indexed by a reduced conflict adaptation effect (i.e., a smaller modulation of reaction times as a function of conflict level) suggesting impaired dynamic adjustment of cognitive control. No significant age-related group differences were observed. Full results are reported in Table S1.

### Fluidity and Excitability differences

Significant group (χ^2^ = 12.38; *p < .001*) and age (χ^2^ = 19.01; *p < .001*) main effects emerged for FI. These results evidence a larger variability of global network stability in ADHD as compared to TD individuals (estimate = 0.38; SE = 0.11; *p = <.001*), and that this index is positively related to the participants’ age. Relevantly, this fluidity difference was independent from frequency bands (Frequency factor: χ2= 1.34; p = .86) (see Fig.2a), implying that these age and group effects occur similarly across frequency bands.

No significant EI group differences emerged (χ2 = 1.9; p = .16). However, we found a significant Group x Region interaction effect on the EI (χ2 = 99.3; *p = .006*). Specifically, the ADHD group was characterized by a larger excitability mainly focused in the frontal areas of the brain, including the dorsolateral prefrontal cortex, frontal gyrus and post-central gyrus, alongside occipital and temporal regions (Fig. 2b).

**Figure 2.**
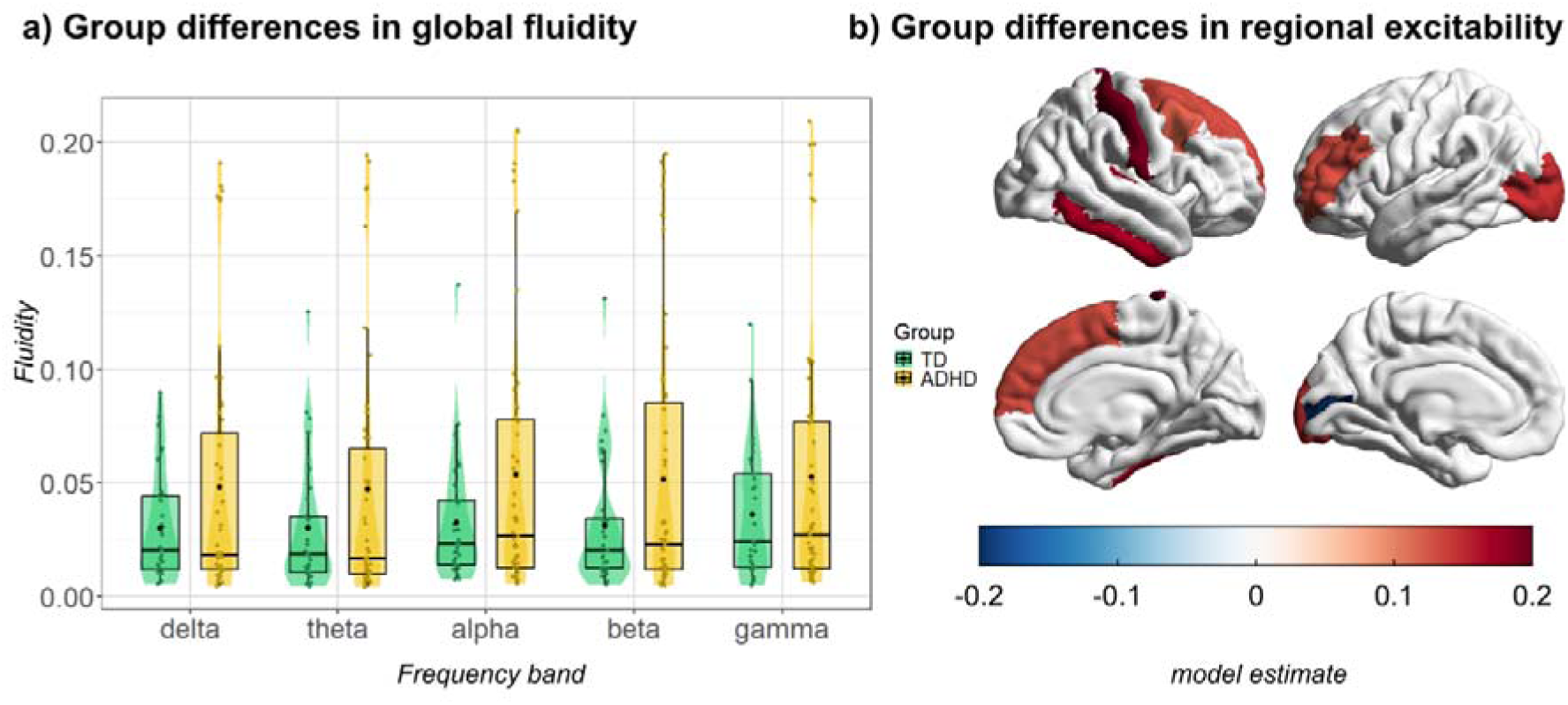
Group difference in Fluidity and Excitability index. Panel (a) shows the group differences in fluidity (y-axis) across the five canonical frequency bands (x-axis) between ADHD (yellow) and TD groups (green). Panel (b) shows the spatial distribution of group differences in cortical excitability between ADHD and TD groups, mapped onto the cortical surface using the Desikan–Killiany (DK) atlas (Desikan et al., 2006). Color values represent β estimates (on a log scale) with red indicating increased excitability in ADHD relative to TD participants and blue indicating decreased excitability. Color scaling is fixed to the range [−0.2, 0.2], and only regions showing statistically significant effects are displayed.

As for the EI-FI interplay, we found (χ2 = 15.35; *p < .001;* See also Table S2) region-specific effects despite no group-level differences (χ2 = 1.59; *p = .17*). Specifically, higher local excitability in frontal (i.e., cingulate and dorsolateral cortices) and occipital (i.e., cuneus) regions is associated with lower fluidity. On the other hand, higher EI values in the occipital cortex, temporal gyrus, intraparietal sulcus, as well as motor and somatosensory cortex, were associated with larger fluidity. These effects were independent from frequency bands (see Fig.3).

**Figure 3.**
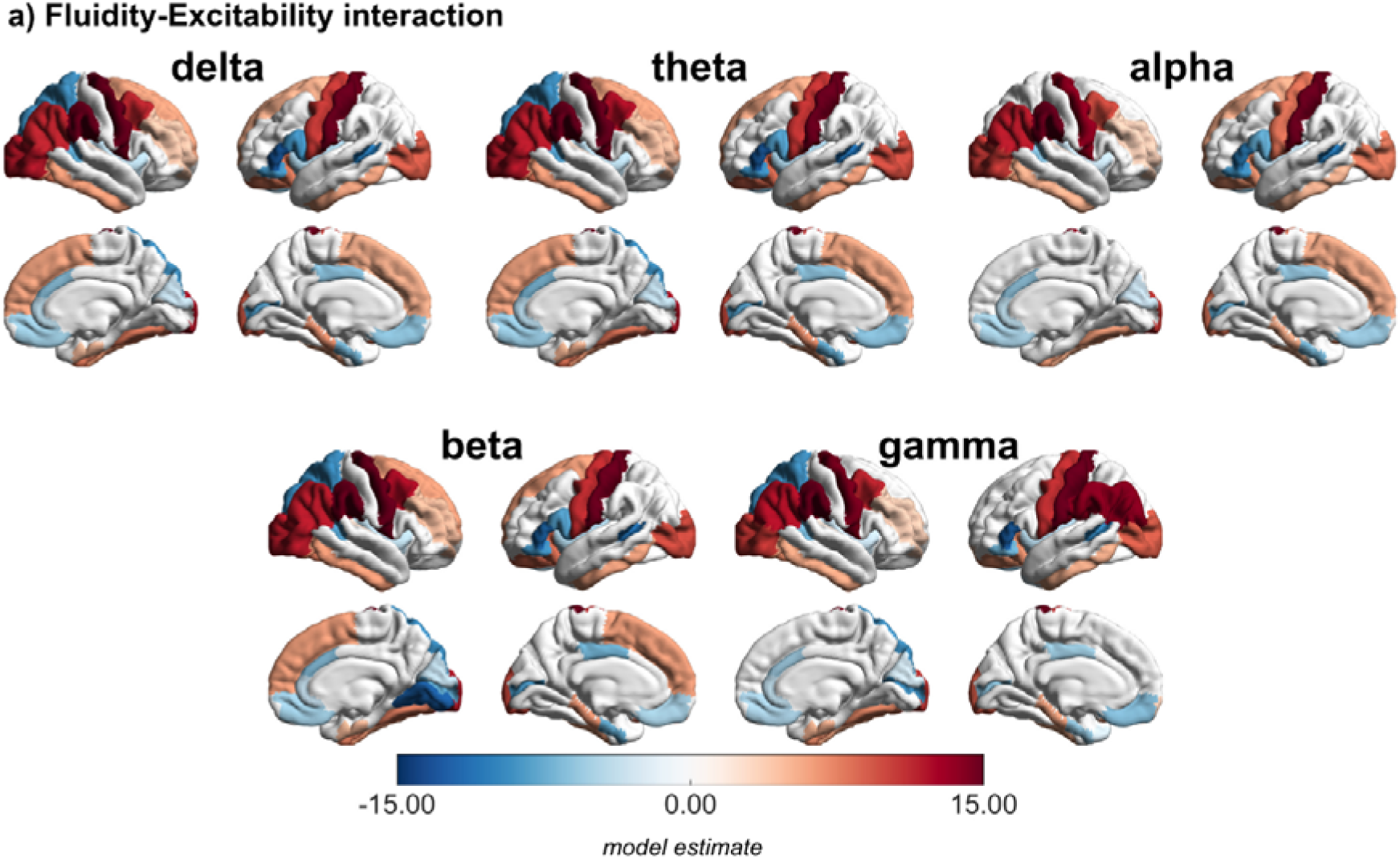
Fluidity-Excitability interaction. Across-group spatial distribution of the interaction between cortical excitability and fluidity across frequency bands. Color values represent β estimates from the linear model, quantifying the strength and direction of the fluidity–excitability interaction within each cortical region. Red colors indicate a positive interaction (greater excitability associated with increased fluidity), whereas blue colors indicate a negative interaction. Values are derived from parcel-wise estimates and projected onto the cortical surface for visualization.

### Brain-Behaviour Interaction

Cross-validation identified an optimal model configuration consisting of two latent components, yielding a significant overall fit (R^2^Y = 17.3% of explained variance, *p* = .011). Further decomposition of the model revealed that the second component was the primary driver of brain-behaviour association, independently explaining 11.8% of the variance (*p* < .05), whereas the first component did not reach significance (see Supporting Information). Investigation of the variable loadings for this significant component highlighted specific contributions from both the behavioural and neural domains. Loadings on the second latent variable (LV2) indicated that, at the behavioural level, the component was characterized by significant contributions (derived from the permutation approach) from Positive Affect (PANAS-P), NEPSY-II Inhibition time scores, phonemic fluency, and participant age (see Fig. 4a; significance assessed via permutation testing). At the neural level, LV2 loadings reflected significant contributions from fluidity within the alpha, beta, and gamma frequency bands. In addition, regional EI loadings on LV2 were prominent in the dorsolateral prefrontal, orbitofrontal, and cingulate cortices, as well as in the cuneus, lingual gyrus, postcentral gyrus, and supramarginal gyrus (see Fig. 4b). Detailed loadings and corresponding p-values for all variables are reported in Tables S3 and S4.

**Figure 4.**
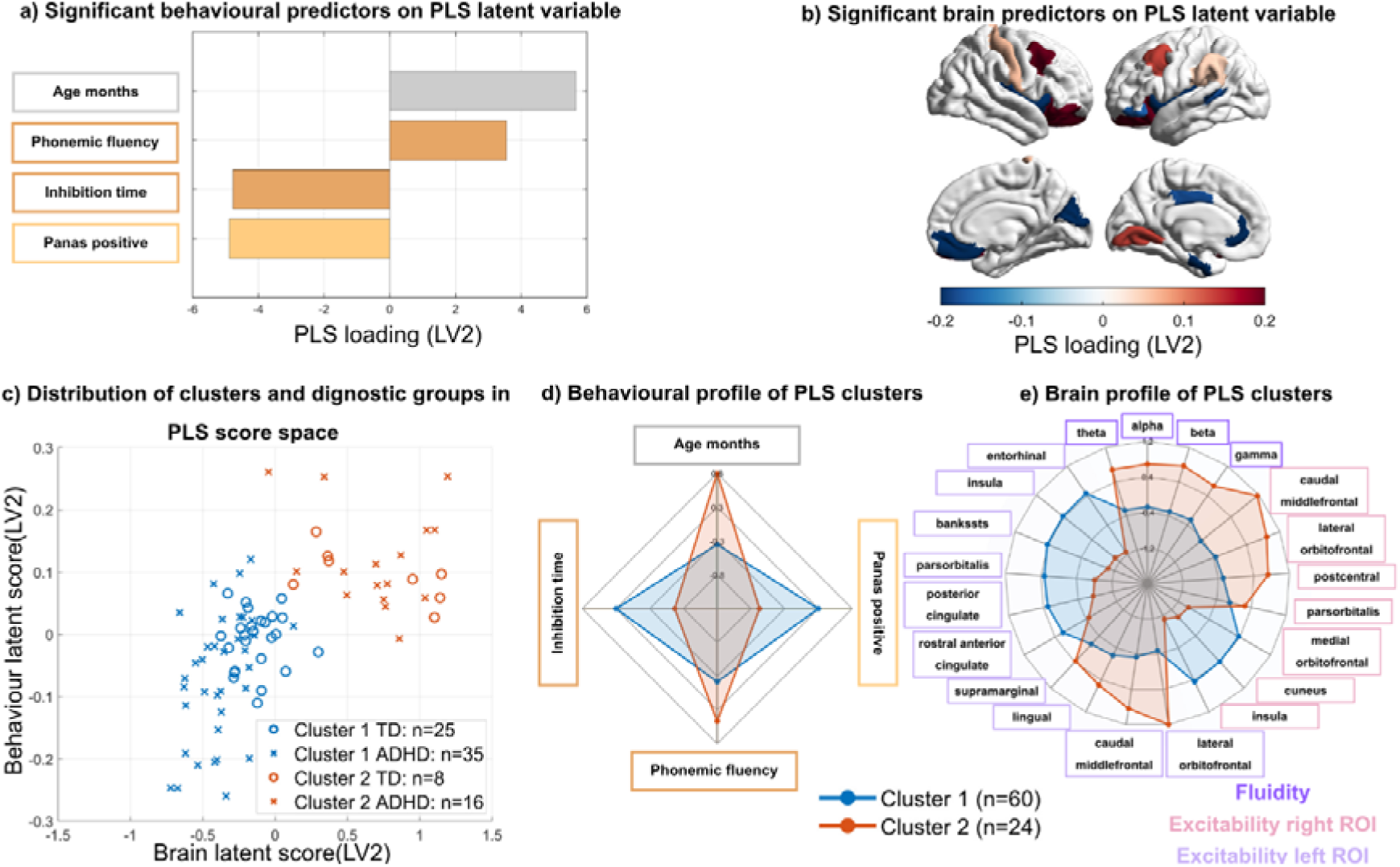
Multimodal characterization of PLS latent variable and related cluster profiles. Panel **(a)** Significant behavioural predictors associated with latent variable 2. Horizontal bars represent behavioural loadings for predictors surviving the significance threshold (*p* < 0.05). **(b)** Significant brain predictors contributing to PLS latent variable 2, displayed as cortical loadings projected onto the DK brain surface. Only regions surviving the significance threshold (*p* < 0.05) are shown, with red and blue colors indicating positive and negative loadings, respectively. **(c)** Distribution of participants in the latent score space defined by brain and behaviour-related scores for latent variable 2. Individuals are shown according to cluster membership (red associated with cluster 1 and blue associated with cluster 2) and diagnostic group (dots associated with the control group and x associated with the ADHD group), allowing visualization of the separation between clusters and clinical groups in the joint PLS space. **(d)** Behavioural profile of the identified PLS clusters. Radar plots summarize cluster-wise mean values for the significant behavioural predictors contributing to latent variable 2. **(e)** Brain profile of the identified PLS clusters. Radar plots summarize cluster-wise mean values for the significant brain features contributing to latent variable 2, including spectral and hemispheric measures.

To further characterize the multivariate structure underlying the significant second component, we applied an unsupervised clustering procedure to participants’ scores on LV2. This step aimed to decompose the global brain–behavior association captured by the component into neurocognitive profiles. The optimal number of clusters *k* was chosen as the one that maximized the mean silhouette value, indicating the best separation and cohesion of clusters. In this case, clustering was performed using k-means with k = 2, selected based on the silhouette index (0.73). Noteworthy, the group was not significantly associated with cluster membership (χ2 = .50, p = .48; see Fig. 4c), suggesting that the identified profiles cut across individuals rather than reflecting group-specific effects (Further details on the clusters sample are reported in Table S5). The two clusters exhibited clearly differentiated patterns across both behavioural and neural dimensions (Full reports in Table S6). Cluster 1 was characterized by younger age, lower phonemic fluency scores, less efficient inhibitory control (longer NEPSY inhibition completion time), and reduced positive affect compared with Cluster 2 (see Fig.4d).

At the neural level, Cluster 1 showed higher excitability across several cortical regions, including the insular, entorhinal, orbitofrontal, and cingulate cortices, as well as temporal areas. In addition, this cluster was associated with reduced network fluidity in all bands except delta. Together, these neural characteristics suggest a profile marked by heightened regional excitability alongside reduced dynamic flexibility of large-scale brain networks (see Fig.4e).

## Discussion

### Behavioural sensitivity: questionnaires outperform performance-based measures

Understanding how intrinsic neurofunctional organization relates to cognitive and behavioural phenotype in ADHD remains a key challenge in clinical developmental neuroscience. Rather than relying on disorder-specific biomarkers, current evidence supports dimensional and developmental frameworks of brain–behaviour interplay. Here, we built on this line of research by combining hdEEG-derived metrics of cortical excitability and large-scale network dynamics with behavioural, cognitive, and emotional dimensions to examine how intrinsic neural activity relates to phenotypic variability of ADHD.

Firstly, we found that children and adolescents with an ADHD diagnosis can be clearly distinguished from TD peers based on behavioural measures derived from parent-report questionnaires assessing attentional, executive, social, affective, and self-regulation domains. By contrast, only a subset of neuropsychological performance-based measures assessing cognitive flexibility and interference control were sufficiently sensitive to significantly differentiate the groups. Specifically, the main group differences emerged in the percentage of correct responses on the WCST and in block-wise conflict adaptation in the Flanker task. Individuals with ADHD showed reduced performance on the WCST, indicating less efficient set-shifting based on explicit rules. In the Flanker task, they exhibited diminished adaptation across high- and low-conflict blocks, as reflected by a smaller modulation of reaction times as a function of conflict level, suggesting a reduced capacity for implicit (i.e., automatic and experience-driven) adjustment to contextual demands.

Taken together, these findings indicate that individuals with ADHD are less efficient in implementing flexible cognition, both when adapting behaviour based on explicit rules and when relying on more implicit, context-sensitive adjustments. This convergence points to a potential common underlying factor related to the flexible deployment of cognitive control, operating across both explicit and implicit modes of regulation.

Overall, behavioural findings align with prior evidence indicating that parent-report questionnaires are more sensitive than performance-based measures in detecting ADHD symptoms. They capture behaviors in everyday contexts, such as inattention, impulsivity, and emotional dysregulation, rather than relying on task performance, which may be influenced by compensatory strategies and individual variability (Miranda et al., 2015; Peterson et al., 2024; Toplak et al., 2013).

### Increase of cortical excitability and network fluidity in ADHD

At the neurofunctional level, ADHD was associated with altered patterns in both local cortical excitability and large-scale network dynamics. Specifically, increased cortical excitability was primarily observed across different regions implicated in CC and attentional regulation, including dorsolateral prefrontal cortex, frontal and post-central gyrus together with occipital and temporal regions. This heightened excitability may reflect a shift from baseline neural states within networks supporting goal-directed behavior, potentially contributing to difficulties in sustaining attention and regulating actions (Castellanos & Proal, 2012; Cortese et al., 2021). In parallel, we observed increased global variability in large-scale brain dynamics (fluidity) in ADHD. While a certain degree of flexibility is essential for adaptive behaviour, excessive variability may compromise the stability required for some cognitive functions, such as sustained attention and CC in general (Armbruster-Genç et al., 2016; Dinstein et al., 2015). Greater fluidity may therefore indicate that networks transit too rapidly between functional states, reducing the persistence of task-relevant configurations (Alderson et al., 2020; Gao et al., 2025). Notably, these differences were independent of the frequency bands, suggesting a general property of neural dynamics rather than an oscillatory-specific effect.

### Region-specific preservation of excitability–network dynamics coupling

Despite these group differences, the relationship between cortical excitability and network dynamics appeared largely preserved across groups. Indeed, regional excitability significantly predicted network fluidity, but this association was expressed across participants instead of differentiating groups. Notably, a region-specific pattern emerged: mesial regions, including the cingulate cortex and cuneus, showed a negative EI-FI interplay. By contrast, dorsal fronto-parietal and temporal regions showed a positive EI-FI association. This pattern is suggestive of a functional dissociation, whereby regions showing negative or positive EI–FI relationships may reflect distinct modes of large-scale network organization. Specifically, this dissociation may be reminiscent of a differential functional interplay between task-negative (default mode) and task-positive (e.g., central executive) networks. We may speculate that this bidirectional organization may suggest that excitability influences network stability differently depending on the functional role of that specific network. In regions compatible with task-negative circuits, increased excitability may stabilize intrinsic activity, reducing fluidity. In contrast, in regions involved in task-positive networks, it may facilitate flexible reconfiguration, increasing fluidity. However, the comparable expression of this coupling across groups indicates that the relationship between local excitability and large-scale dynamics reflects a general organizational principle of brain function instead of a core ADHD neural fingerprint.

### A dimensional neurodevelopmental organization of brain–behaviour relationships

When neural and behavioral measures were combined within a multivariate approach, a latent brain–behavior dimension emerged linking excitability and fluidity with age, inhibitory control, verbal fluency, and state-dependent positive affect. The contributing regions included nodes from attentional and CC networks, as well as default mode areas, supporting the balance between internally and externally oriented cognition (Weber et al., 2022). Variability in these systems may influence the allocation of attentional resources and the regulation of goal-directed behavior. Neural configurations characterized by stable control networks and appropriate modulation of default mode activity may support more efficient EF, strategic vocabulary access, and emotional regulation. The emergence of age as a significant variable in explaining brain-behaviour axis highlights the developmental nature of this interplay suggesting that neural and behavioral maturation follow shared trajectories.

Importantly, these multivariate patterns did not map onto groups. Clustering analyses identified two distinct profiles driven primarily by age and neurocognitive differences rather than by diagnosis. This finding may support a dimensional perspective in which ADHD reflects a shifted position along broader neurodevelopmental continua rather than a discrete category. Consistent with transdiagnostic frameworks, variation in neurofunctional organization appears better captured along continuous dimensions than by disorder-specific biomarkers (Astle et al., 2022; Chopra et al., 2025; Hoy et al., 2023).

From a clinical perspective, these findings suggest that measures of cortical excitability and network dynamics could help identify individual neurocognitive fingerprints that go beyond traditional diagnostic labels, potentially informing individualized interventions targeting and/or monitoring network stability or excitability.

In conclusion, our study shows that ADHD is associated with increased cortical excitability and greater variability in large-scale neural dynamics. However, brain–behavior relationships unfold along developmental dimensions that transcend diagnostic categories, and everyday behavioral measures remain the most sensitive indicators of ADHD-related differences. Overall, these findings support a dimensional, developmentally grounded framework that emphasizes integrated neurofunctional profiles over categorical biomarkers.

## Conclusion

ADHD shows increased cortical excitability and more variable brain dynamics, but these differences do not define a distinct neural category. Instead, brain–behaviour relationships follow a shared developmental dimension across individuals. This supports a dimensional, transdiagnostic view of neurodevelopment beyond diagnostic labels

## Fundings

The study was funded by: Ricerca Corrente 2026 from Italian Ministry of Health; Complementary National Plan PNC-I.1 “Research initiatives for 684 innovative technologies and pathways in the health and welfare sector” D.D. 931 of 06/06/2022, DARE - 685 DigitAl lifelong pRevEntion initiative, code PNC0000002, CUP: B53C22006450001; Italian Ministry of University and Research under the National Recovery and Resilience Plan (PNRR), funded by the European Union – NextGenerationEU. Project ID:C96E23000850001.

## Acknowledgements

The authors wish to thank all the families who participated in this study. Additional thanks to the students who contributed to data collection: C. Cagnucci, M. D’Angelo, I. Esposito, L. Iannazzo, G. Zampieri, E. R. Zacheo. We would like to especially thank the local ADHD family associations “Zuppa di Sasso”, “Gemelli ADHD”, and “Stelle sulla Terra” for their support in participant recruitment. A special Thanks to Prof. Z Alex, for teaching us never to give up, even when everything seems lost.

## Ethical Information

All parents provided written consent. All experimental procedures were approved by the Ethics Committee of the School of Psychology of the University of Padua (protocol no. 387-a) and from CET Area Nord Veneto (N. 0012604/24).

## Data availability

All materials and data are available on the Open Science Framework (OSF): https://osf.io/fkhe9/overview

## Key Points What’s known?

- ADHD shows heterogeneous cognitive and neural profiles, with no reliable disorder-specific biomarker, supporting dimensional models over categorical diagnoses.

## What’s new?

- Using hdEEG, we show increased brain fluidity and cortical excitability in ADHD, alongside a region-specific excitability–fluidity coupling preserved across groups.
- A latent brain–behaviour dimension links neural dynamics with age, inhibitory control, verbal fluency, and positive affect, without separating ADHD from typically developing peers.

## What’s relevant?

- Findings support a developmental, dimensional framework where neurofunctional variation reflects continuous dimensions, informing more precise, individualized approaches beyond diagnostic labels.

## Notes

https://osf.io/fkhe9/overview

### Competing Interest Statement

The authors have declared no competing interest.

