## Supporting Materials for "Where is ADHD in the brain? Evidence for a neurodevelopmental continuum of brain dynamics"

**Participants Details**

**Inclusion and exclusion criteria.** Fluid intelligence was assessed using the Coloured Progressive Matrices (CPM; Belacchi et al., 2008; Raven et al., 1998) and the Standard Progressive Matrices (SPM; Raven et al., 2000), and one participant was excluded for scoring at least two standard deviations below the population mean on either measure. An additional participant was excluded due to poor-quality resting-state hdEEG data caused by excessive noise or movement artefacts. Participants with known diagnosis of sensory, neurological, or neuropsychiatric disorders and without normal or corrected-to-normal vision were also excluded. Further exclusion criteria included insufficient performance in the experimental task, defined as overall accuracy below 60% or block-level accuracy below 50% (n = 3). Finally, ten participants were excluded due to incomplete neuropsychological assessments or missing parent-reported questionnaires.

**Pharmacological characteristics and subtypes.** Within the ADHD group: 35.29% of participants were unmedicated, 41.18% were treated with methylphenidate, 11.76% received a combination of methylphenidate and risperidone, 3.92% treated with risperidone only, 1.96% with SSRIs, 1.96% a combination of SSRIs and risperidone, and 3.92% with all three classes of psychotropic medication. Regarding ADHD subtypes, 37.25% were classified as combined type, 9.80% as inattentive, and only 1.96% as hyperactive/impulsive, while subtype information was not specified for 50.98% of the sample.

**Behavioral and neuropsychological Assessment**

Participants were administered a task aimed at assessing cognitive flexibility, namely the Wisconsin Card Sorting Test (WCST; Heaton, 1981), in its computerized 64-card version implemented using the PEBL software (version 2.1; Mueller, 2011). In this version, four stimulus cards are presented on the screen, and on each trial, a response card appears. The stimulus cards differ in color, shape, and number of items. Participants are required to match each response card to one of the stimulus cards by inferring the underlying sorting rule, which is never explicitly stated but can be deduced from feedback. As the sorting rule changes multiple times during the task, participants must flexibly update previously learned strategies and adapt their responses accordingly.

Subsequently, participants completed paper-and-pencil tasks. Short-term memory was assessed using the Backward Digit Span subtest from the Neuropsychological Assessment Battery for Developmental Age (BVN 5–11; Bisiacchi et al., 2005; BVN 12–18; Gugliotta, 2009). From the same battery, a phonemic fluency task was administered to assess lexical access speed and impulsivity. Finally, inhibitory control was further examined using the Inhibition subtest of the NEPSY-II battery (Urgesi, Campanella, & Fabbro, 2011). This task includes three conditions (A, B, C), each probing different executive processes; the condition of interest for the present study was condition B (inhibition), in which participants are required to provide a response opposite to the presented stimulus as quickly as possible.

Participants also completed two self-report questionnaires: the PANAS-C short form (Ciucci et al., 2017; Ebesutani et al., 2012; Thompson, 2007) and the Body-Map. The PANAS-C consists of two mood-related subscales, each comprising five items: one assessing positive affect and the other negative affect. Participants are asked to rate, on a scale from 1 (“very slightly or not at all”) to 5 (“extremely”), the extent to which each listed emotional state reflects their experience. The Body-Map is a scale designed to assess participants’ capacity for interoceptive awareness. It includes eight different body areas and requires participants to indicate, on a scale from 1 (“not at all”) to 7 (“very much”), the extent to which they perceive and recognise emotions in each area.

All participants also completed a modified version of the Flanker task (Gonthier et al., 2021; Toffoli et al. 2025), designed to assess the ability to manage cognitive interference under contexts that differentially facilitate or hinder performance. Specifically, the task manipulated contextual demands by varying the proportion of trials’ congruency: mostly incongruent (MI; higher frequency of distractors), mostly congruent (MC; lower frequency of distractors), and non-predictive (NP; 50% congruent and incongruent trials). In each trial, a horizontal array of fish stimuli pointing either left or right was presented. Participants were instructed to indicate the direction of the central fish as quickly as possible using a keyboard, allowing assessment of both reaction times (RTs) and accuracy. The total duration of the task was approximately 20 minutes, including breaks. Before the experimental session, a practice block was administered to ensure that participants understood the instructions.

Parents completed the questionnaires through the Qualtrics Survey platform (Qualtrics, 2019).
The Behavior Rating Inventory of Executive Function – Second Edition (BRIEF-2; Gioia et al., 2002; Italian adaptation by Marano et al., 2016) is a questionnaire designed to assess executive functioning in everyday contexts, thus providing an ecologically valid measure of these abilities. It allows the investigation of eight specific domains (inhibition, emotional control, flexibility, working memory, planning/organization, monitoring, initiation, and task monitoring), which are grouped into three main indices: the Behavioral Regulation Index (BRI), the Emotional Regulation Index (ERI), and the Cognitive Regulation Index (CRI), as well as a Global Executive Composite (GEC).
The Conners’ Parent Rating Scale – Revised (CPRS-R; Conners et al., 1998; Italian adaptation by Nobile et al., 2007) specifically assesses core ADHD symptoms and associated behavioural problems. It comprises several subscales (Inattention/Cognitive Problems, Hyperactivity, Anxiety/Shyness, Perfectionism, Social Problems, Psychosomatic Problems, Restlessness/Impulsivity, and Emotional Lability), reflecting the main symptomatic and functional domains associated with the disorder. It also includes global indices such as the Conners’ Global Index and DSM-oriented scales. The Social Responsiveness Scale – Second Edition (SRS-2; Italian adaptation by D’Ardia et al., 2021) assesses the quality of social interactions, reciprocal communication, social motivation, and the presence of stereotyped behaviours, and was used as a screening tool to evaluate traits associated with autism spectrum disorder. It provides two main composite indices: Social Communication and Interaction (SCI) and Restricted and Repetitive Behaviors (RRB).

**Fluidity Fomulas**

*[Formula 1]* $dFC(t\_1,t\_2) = corr[UpperTri(FC(t\_1),UpperTri(FC(t\_2 )]$

*[Formula 2]* $dFC\left( t_{1},t_{2} \right)\in R^{N x N}, for M time windows$

*[Formula 3]* $fluidity= Var\left[ UpperTri\left( dFC - B \right) \right]$

*[Formula 4]* $Bij= 1 if\left[ t_{i}-\frac{t_{win}}{2}; t_{i}+\frac{t_{win}}{2} \right]\cap\left[ t_{j}-\frac{t_{win}}{2}; t_{j}+\frac{t_{win}}{2} \right] \neq\emptyset; 0 otherwise$

*[Formula 5]* $offset=\frac{n_{overlap}}{t_{win}- n_{overlap}}+1$

**Additional PLS Results**

Permutation testing revealed that the first latent variable (LV1) was not statistically significant (5.5% explained variance, p = .244), indicating that it does not reliably capture brain–behaviour covariance. In contrast, the second latent variable (LV2) was statistically significant (11.8% explained variance, p < .05) and was therefore retained for interpretation in the main analyses.

|  | Variables | Mean ± SD Controls | Mean ± SD ADHD | t | p-value (FDR corrected) |  |
| --- | --- | --- | --- | --- | --- | --- |
|  | Age (in months) | 140.58 ± 29.08 | 152.75 ± 27.7 | -1.91 | 0.16 |  |
| Self-report Questionnaires | Panas Positive | 14.76 ± 4.47 | 15.10 ± 5.00 | -0.33 | 0.84 |  |
|  | Panas Negative | 5.85 ± 1.15 | 6.37 ± 2.24 | -1.41 | 0.33 |  |
|  | Body-Map | 30.79 ± 6.93 | 28.73 ± 8.22 | 1.24 | 0.39 |  |
| Parent-report Questionnaires | Conners  (Oppositivity) | 6.27 ± 4.75 | 14.67 ± 6.76 | -6.68 | <0.001 | * |
|  | Conners (Cognitive problems and disattention) | 5.39 ± 4.93 | 20.78 ± 7.26 | -11.57 | <0.001 | * |
|  | Conners  (Hyperactivity) | 2.33 ± 3.33 | 12.10 ± 5.77 | -9.82 | <0.001 | * |
|  | Conners  (Anxiety and shyness) | 3.33 ± 3.41 | 6.73 ± 4.99 | -3.70 | <0.001 | * |
|  | Conners (Perfectionism) | 2.52 ± 2.40 | 6.02 ± 4.08 | -4.95 | <0.001 | * |
|  | Conners  (Social problems) | 1.12 ± 1.93 | 4.02 ± 3.64 | -4.75 | <0.001 | * |
|  | Conners (Psicosomatic problems) | 1.52 ± 1.94 | 3.57 ± 2.67 | -4.08 | <0.001 | * |
|  | Conners  (ADHD index) | 5.03 ± 4.93 | 22.61 ± 7.27 | -13.21 | <0.001 | * |
|  | Conners  (Restlessness/ Impulsivity) | 2.67 ± 2.75 | 11.92 ± 4.35 | -11.95 | <0.001 | * |
|  | Conners  (Emotional Lability) | 0.85 ± 1.37 | 2.76 ± 2.15 | -4.99 | <0.001 | * |
|  | Conners  (Global index) | 3.52 ± 3.69 | 14.69 ± 5.49 | -11.14 | <0.001 | * |
|  | Conners  (DSM Inattention) | 4.91 ± 4.30 | 16.71 ± 6.26 | -10.23 | <0.001 | * |
|  | Conners  (DSM Hyperactivity/  Impulsivity) | 4.24 ± 3.66 | 14.10 ± 5.66 | -9.69 | <0.001 | * |
|  | Conners (DSM-IV total scale) | 8.21 ± 6.80 | 30.12 ± 10.9 | -11.34 | <0.001 | * |
|  | Brief  (Self-Monitoring) | 6.30 ± 1.74 | 9.41 ± 2.09 | -7.38 | <0.001 | * |
|  | Brief  (Inhibition) | 10.61 ± 2.89 | 16.49 ± 3.06 | -8.90 | <0.001 | * |
|  | Brief  (Behavioral Regulation Index BRI) | 16.91 ± 4.10 | 25.71 ± 4.60 | -9.15 | <0.001 | * |
|  | Brief  (Shift) | 10.36 ± 2.03 | 15.92 ± 3.42 | -9.35 | <0.001 | * |
|  | Brief  (Emotional Control) | 11.15 ± 2.64 | 15.96 ± 3.64 | -7.01 | <0.001 | * |
|  | Brief  (Emotion Regulation Index ERI) | 21.52 ± 3.88 | 31.86 ± 6.23 | -9.37 | <0.001 | * |
|  | Brief  (Initiate) | 6.55 ± 1.30 | 10.33 ± 2.16 | -10.02 | <0.001 | * |
|  | Brief (Working memory) | 10.52 ± 2.35 | 17.59 ± 3.23 | -11.60 | <0.001 | * |
|  | Brief  (Planning/Organization) | 11.06 ± 2.74 | 18.14 ± 3.62 | -10.18 | <0.001 | * |
|  | Brief  (Task monitoring) | 7.85 ± 2.18 | 11.18 ± 2.16 | -6.85 | <0.001 | * |
|  | Brief  (Organization of materials) | 8.30 ± 1.98 | 11.53 ± 2.93 | -6.03 | <0.001 | * |
|  | Brief  (Cognitive Regulation Index CRI) | 44.27 ± 8.86 | 69.06 ± 11.37 | -11.18 | <0.001 | * |
|  | Brief  (Global Executive Composite GEC) | 82.70 ± 13.77 | 126.63 ± 17.26 | -12.91 | <0.001 | * |
|  | SRS  (Social awareness AWR) | 5.82 ± 2.34 | 9.43 ± 3.24 | -5.93 | <0.001 | * |
|  | SRS  (Social cognition COG) | 4.91 ± 2.42 | 14.04 ± 5.35 | -10.63 | <0.001 | * |
|  | SRS  (Social communication COM) | 7.85 ± 4.92 | 21.75 ± 10.24 | -8.32 | <0.001 | * |
|  | SRS  (Social motivation MOT) | 3.64 ± 2.55 | 9.69 ± 6.14 | -6.25 | <0.001 | * |
|  | SRS  (Restricted Interests and Repetitive Behavior RRB) | 3.09 ± 2.85 | 12.18 ± 5.99 | -9.32 | <0.001 | * |
|  | SRS  (Social Communication and Interaction SCI) | 22.21 ± 9.96 | 54.90 ± 22.45 | -9.11 | <0.001 | * |
| Neuro- psychological measures | Backward digit span | 4.00 ± 1.03 | 3.75 ± 1.25 | 1.02 | 0.48 |  |
|  | Inhibition errors NEPSY | 3.64 ± 3.25 | 3.22 ± 3.98 | 0.53 | 0.73 |  |
|  | Inhibition time NEPSY | 62.94 ± 14.43 | 60.40 ± 14.03 | 0.80 | 0.59 |  |
|  | Phonemic fluency BVN | 32.03 ± 10.14 | 29.98 ± 9.34 | 0.93 | 0.53 |  |
|  | WCST  (Correct percent) | 79.31 ± 9.02 | 72.11 ± 12.32 | 3.09 | 0.01 | * |
|  | WCST  (Learning to learn index) | 1.76 ± 4.34 | 0.96 ± 3.09 | 0.91 | 0.54 |  |
|  | WCST  (Perseverative  errors percent) | 11.08 ± 4.76 | 12.12 ± 6.89 | -0.82 | 0.58 |  |
|  | WCST  (CLR percent) | 71.88 ± 12.65 | 62.41 ± 17.39 | 2.88 | 0.02 | * |
| Modified Flanker task | Flanker effects accuracy | 0.07 ± 0.07 | 0.09 ± 0.07 | -1.22 | 0.39 |  |
|  | Flanker effects  RTs | -65.93 ± 24.45 | -73.65 ± 23.40 | 1.44 | 0.32 |  |
|  | Delta  (Flanker effects accuracy – MC vs MI) | 0.03 ± 0.07 | 0.05 ± 0.08 | -1.30 | 0.37 |  |
|  | Delta  (Flanker effects  RTs – MC vs MI) | -18.57 ± 29.84 | -38.98 ± 46.82 | 2.44 | 0.05 |  |
|  | Delta  (Flanker effects accuracy – MC vs NP) | 0.02 ± 0.10 | 0.07 ± 0.10 | -1.91 | 0.16 |  |
|  | Delta  (Flanker effects  RTs – MC vs NP) | -3.97 ± 47.11 | -32.59 ± 47.56 | 2.71 | 0.03 | * |
|  | Delta  (Flanker effects accuracy – MI vs NP) | -0.01 ± 0.08 | 0.01 ± 0.09 | -0.98 | 0.50 |  |
|  | Delta  (Flanker effects  RTs – MI vs NP) | 14.61 ± 44.63 | 6.39 ± 45.20 | 0.82 | 0.58 |  |
|  | Delta  (Incongruent effects accuracy – MC vs MI) | -0.02 ± 0.08 | -0.04 ± 0.08 | 0.84 | 0.58 |  |
|  | Delta  (Incongruent effects  RTs – MC vs MI) | -20.95 ± 37.08 | 3.93 ± 41.57 | -2.86 | 0.02 | * |
|  | Delta  (Incongruent effects accuracy – MC vs NP) | -0.02 ± 0.09 | -0.05 ± 0.09 | 1.71 | 0.22 |  |
|  | Delta  (Incongruent effects  RTs – MC vs NP) | -19.30 ± 53.28 | 1.48 ± 44.97 | -1.85 | 0.17 |  |
|  | Delta  (Incongruent effects accuracy – MI vs NP) | 0.01 ± 0.09 | -0.01 ± 0.07 | 1.04 | 0.48 |  |
|  | Delta  (Incongruent effects  RTs – MI vs NP) | 1.65 ± 36.54 | -2.45 ± 37.80 | 0.50 | 0.75 |  |

**Table S1.** Group differences in neural and behavioural variables (FDR-corrected p-values). Independent-samples t-tests were performed across groups for all neural and behavioural variables. Measures were grouped into macro-domains (questionnaires, neuropsychological measures, and Flanker indices). The table reports t-values, mean ± SD for each group, and FDR-corrected p-values.

| Regions | Bands | Estimate | p-value (FDR corrected) |
| --- | --- | --- | --- |
| Bankssts L | alpha | -10.32 | <0.001 |
| Bankssts L | beta | -11.01 | <0.001 |
| Bankssts L | delta | -10.38 | <0.001 |
| Bankssts L | gamma | -9.60 | <0.001 |
| Bankssts L | theta | -10.91 | <0.001 |
| Bankssts R | alpha | -6.54 | <0.01 |
| Bankssts R | beta | -8.28 | <0.01 |
| Bankssts R | delta | -7.90 | <0.01 |
| Bankssts R | gamma | -6.96 | <0.01 |
| Bankssts R | theta | -8.35 | <0.01 |
| Caudal anterior cingulate R | alpha | -5.95 | <0.001 |
| Caudal anterior cingulate R | beta | -6.38 | <0.001 |
| Caudal anterior cingulate R | delta | -6.162 | <0.001 |
| Caudal anterior cingulate R | gamma | -5.40 | <0.001 |
| Caudal anterior cingulate R | theta | -6.73 | <0.001 |
| Caudal middle frontal R | alpha | 9.80 | <0.01 |
| Caudal middle frontal R | beta | 12.20 | <0.001 |
| Caudal middle frontal R | delta | 11.36 | <0.001 |
| Caudal middle frontal R | gamma | 11.40 | <0.001 |
| Caudal middle frontal R | theta | 12.14 | <0.001 |
| Cuneus R | alpha | -3.31 | <0.05 |
| Cuneus R | beta | -3.84 | <0.01 |
| Cuneus R | delta | -3.56 | <0.01 |
| Cuneus R | gamma | -2.83 | <0.05 |
| Cuneus R | theta | -3.76 | <0.01 |
| Entorhinal L | alpha | -6.80 | <0.001 |
| Entorhinal L | beta | -7.58 | <0.001 |
| Entorhinal L | delta | -6.86 | <0.001 |
| Entorhinal L | gamma | -5.76 | <0.01 |
| Entorhinal L | theta | -7.60 | <0.001 |
| Entorhinal R | beta | 4.74 | <0.05 |
| Entorhinal R | delta | 4.67 | <0.05 |
| Entorhinal R | gamma | 5.13 | <0.05 |
| Entorhinal R | theta | 5.00 | <0.05 |
| Frontalpole L | gamma | 3.19 | <0.05 |
| Fusiform R | alpha | 6.24 | <0.05 |
| Fusiform R | beta | 7.47 | <0.01 |
| Fusiform R | delta | 7.70 | <0.01 |
| Fusiform R | gamma | 7.15 | <0.05 |
| Fusiform R | theta | 7.97 | <0.01 |
| Inferior parietal L | gamma | 14.02 | <0.05 |
| Inferior parietal R | alpha | 11.73 | <0.05 |
| Inferior parietal R | beta | 11.78 | <0.05 |
| Inferior parietal R | delta | 11.40 | <0.05 |
| Inferior parietal R | gamma | 12.14 | <0.05 |
| Inferior parietal R | theta | 11.79 | <0.05 |
| Inferior temporal L | alpha | 4.96 | <0.05 |
| Inferior temporal L | beta | 5.97 | <0.01 |
| Inferior temporal L | delta | 5.80 | <0.01 |
| Inferior temporal L | gamma | 5.29 | <0.05 |
| Inferior temporal L | theta | 5.99 | <0.01 |
| Inferior temporal R | alpha | 5.29 | <0.001 |
| Inferior temporal R | beta | 6.36 | <0.001 |
| Inferior temporal R | delta | 6.23 | <0.001 |
| Inferior temporal R | gamma | 5.67 | <0.001 |
| Inferior temporal R | theta | 6.32 | <0.001 |
| Insula L | alpha | -3.73 | <0.01 |
| Insula L | beta | -4.69 | <0.001 |
| Insula L | delta | -4.63 | <0.001 |
| Insula L | gamma | -4.43 | <0.001 |
| Insula L | theta | -4.80 | <0.001 |
| Insula R | alpha | -3.46 | <0.01 |
| Insula R | beta | -3.89 | <0.01 |
| Insula R | delta | -4.07 | <0.001 |
| Insula R | gamma | -3.32 | <0.01 |
| Insula R | theta | -4.18 | <0.001 |
| Lateral occipital L | alpha | 7.80 | <0.01 |
| Lateral occipital L | beta | 9.78 | <0.001 |
| Lateral occipital L | delta | 9.08 | <0.001 |
| Lateral occipital L | gamma | 8.91 | <0.001 |
| Lateral occipital L | theta | 9.64 | <0.001 |
| Lateral occipital R | alpha | 10.54 | <0.05 |
| Lateral occipital R | beta | 11.75 | <0.01 |
| Lateral occipital R | delta | 11.87 | <0.01 |
| Lateral occipital R | gamma | 10.59 | <0.05 |
| Lateral occipital R | theta | 11.86 | <0.01 |
| Lateral orbitofrontal L | alpha | 6.67 | <0.01 |
| Lateral orbitofrontal L | beta | 8.09 | <0.001 |
| Lateral orbitofrontal L | delta | 7.58 | <0.001 |
| Lateral orbitofrontal L | gamma | 6.66 | <0.01 |
| Lateral orbitofrontal L | theta | 8.23 | <0.001 |
| Lingual R | beta | -12.15 | <0.05 |
| Medial orbitofrontal L | alpha | -5.53 | <0.01 |
| Medial orbitofrontal L | beta | -5.02 | <0.05 |
| Medial orbitofrontal L | delta | -6.01 | <0.01 |
| Medial orbitofrontal L | gamma | -6.43 | <0.01 |
| Medial orbitofrontal L | theta | -6.23 | <0.01 |
| Medial orbitofrontal R | alpha | -4.93 | <0.01 |
| Medial orbitofrontal R | beta | -5.31 | <0.01 |
| Medial orbitofrontal R | delta | -5.40 | <0.01 |
| Medial orbitofrontal R | gamma | -4.80 | <0.05 |
| Medial orbitofrontal R | theta | -5.72 | <0.01 |
| Parahippocampal L | alpha | 5.39 | <0.01 |
| Parahippocampal L | beta | 6.50 | <0.001 |
| Parahippocampal L | delta | 6.60 | <0.001 |
| Parahippocampal L | gamma | 5.96 | <0.01 |
| Parahippocampal L | theta | 6.80 | <0.001 |
| Pars opercularis L | alpha | -7.64 | <0.05 |
| Pars opercularis L | beta | -8.29 | <0.05 |
| Pars opercularis L | delta | -8.88 | <0.05 |
| Pars opercularis L | theta | -8.24 | <0.05 |
| Pars orbitalis L | alpha | -8.90 | <0.01 |
| Pars orbitalis L | beta | -8.80 | <0.01 |
| Pars orbitalis L | delta | -8.72 | <0.01 |
| Pars orbitalis L | gamma | -7.22 | <0.05 |
| Pars orbitalis L | theta | -9.06 | <0.01 |
| Pars triangularis L | alpha | -10.06 | <0.01 |
| Pars triangularis L | beta | -11.47 | <0.01 |
| Pars triangularis L | delta | -12.28 | <0.001 |
| Pars triangularis L | gamma | -11.93 | <0.01 |
| Pars triangularis L | theta | -12.13 | <0.01 |
| Pericalcarine L | alpha | -7.72 | <0.01 |
| Pericalcarine L | beta | -8.48 | <0.01 |
| Pericalcarine L | delta | -8.31 | <0.001 |
| Pericalcarine L | gamma | -6.06 | <0.05 |
| Pericalcarine L | theta | -8.20 | <0.01 |
| Pericalcarine R | beta | -7.81 | <0.05 |
| Pericalcarine R | gamma | -8.93 | <0.05 |
| Postcentral L | alpha | 14.89 | <0.01 |
| Postcentral L | beta | 18.63 | <0.01 |
| Postcentral L | delta | 18.84 | <0.001 |
| Postcentral L | gamma | 16.78 | <0.01 |
| Postcentral L | theta | 19.24 | <0.001 |
| Posterior cingulate L | alpha | -5.79 | <0.01 |
| Posterior cingulate L | beta | -7.14 | <0.01 |
| Posterior cingulate L | delta | -6.30 | <0.01 |
| Posterior cingulate L | gamma | -5.42 | <0.05 |
| Posterior cingulate L | theta | -6.20 | <0.01 |
| Precentral L | alpha | 8.10 | <0.05 |
| Precentral L | beta | 11.12 | <0.01 |
| Precentral L | delta | 10.46 | <0.01 |
| Precentral L | gamma | 11.01 | <0.01 |
| Precentral L | theta | 10.74 | <0.01 |
| Precentral R | alpha | 13.36 | <0.01 |
| Precentral R | beta | 16.78 | <0.001 |
| Precentral R | delta | 15.49 | <0.001 |
| Precentral R | gamma | 14.39 | <0.01 |
| Precentral R | theta | 16.31 | <0.001 |
| Rostral anterior cingulate R | alpha | -4.22 | <0.01 |
| Rostral anterior cingulate R | beta | -3.75 | <0.05 |
| Rostral anterior cingulate R | delta | -4.11 | <0.01 |
| Rostral anterior cingulate R | gamma | -3.97 | <0.01 |
| Rostral anterior cingulate R | theta | -4.37 | <0.01 |
| Rostral middle frontal R | alpha | 3.45 | <0.01 |
| Rostral middle frontal R | beta | 4.54 | <0.01 |
| Rostral middle frontal R | delta | 3.96 | <0.01 |
| Rostral middle frontal R | gamma | 3.61 | <0.05 |
| Rostral middle frontal R | theta | 4.40 | <0.01 |
| Superior frontal L | alpha | 5.46 | <0.05 |
| Superior frontal L | beta | 6.17 | <0.05 |
| Superior frontal L | delta | 5.32 | <0.05 |
| Superior frontal L | theta | 5.67 | <0.05 |
| Superior frontal R | beta | 6.16 | <0.05 |
| Superior frontal R | delta | 5.89 | <0.05 |
| Superior frontal R | theta | 5.92 | <0.05 |
| Superior parietal R | beta | -9.25 | <0.05 |
| Superior parietal R | delta | -8.91 | <0.05 |
| Superior parietal R | gamma | -8.95 | <0.05 |
| Superior parietal R | theta | -9.08 | <0.05 |
| Supramarginal L | gamma | 13.85 | <0.05 |
| Supramarginal R | alpha | 18.43 | <0.01 |
| Supramarginal R | beta | 22.36 | <0.01 |
| Supramarginal R | delta | 23.34 | <0.001 |
| Supramarginal R | gamma | 25.14 | <0.001 |
| Supramarginal R | theta | 23.19 | <0.001 |
| Temporal pole L | gamma | -2.36 | <0.05 |
| Transverse temporal R | alpha | 1.82 | <0.01 |
| Transverse temporal R | beta | 2.12 | <0.001 |
| Transverse temporal R | delta | 2.15 | <0.001 |
| Transverse temporal R | gamma | 1.65 | <0.01 |
| Transverse temporal R | theta | 2.18 | <0.001 |

**Table S2.** Cortical regions showing a significant main effect of excitability on fluidity. For each region and frequency band, estimates (β) were obtained from a mass-univariate linear model. P-values were corrected for multiple comparisons across regions within each frequency band using false discovery rate (FDR). Only region–band pairs showing a significant main effect of excitability after FDR correction (p < 0.05) are reported.

| **Variable** | **Loading** | **p-value** |
| --- | --- | --- |
| Age (in months) | 5.65 | 0.002 |
| Panas positive | -4.88 | 0.001 |
| Inhibition time NEPSY | -4.77 | 0.009 |
| Phonemic fluency BVN | 3.55 | 0.05 |

**Table S3.** Loadings and permutation-based p-values for behavioural variables contributing significantly to the second latent variable (LV2) identified by the PLS analysis. Behavioural loadings reflect the strength and direction of their contribution to the latent brain–behaviour association. Statistical significance was assessed using permutation testing (10,000 permutations), and only variables with p < .05 are reported.

| **Variable** | **Loading** | **p-value** |
| --- | --- | --- |
| Insula R | -0.53 | 0.01 |
| Insula L | -0.50 | 0.001 |
| Cuneus R | -0.40 | 0.01 |
| Entorhinal L | -0.37 | <0.001 |
| Pars orbitalis R | 0.29 | 0.007 |
| Medial orbito frontal R | -0.30 | 0.006 |
| Lateral orbito frontal L | 0.27 | <0.001 |
| Lateral orbito frontal R | 0.25 | 0.005 |
| Caudal middle frontalR | 0.21 | 0.01 |
| Rostral anterior cingulate L | -0.20 | 0.02 |
| Posterior cingulate L | -0.18 | 0.04 |
| Bankssts L | -0.18 | 0.01 |
| Pars orbitalis L | -0.17 | 0.004 |
| Caudal middle frontal L | 0.13 | 0.01 |
| Lingual L | 0.13 | 0.004 |
| Post central R | 0.06 | 0.02 |
| Supramarginal L | 0.05 | 0.01 |
| Alpha | 0.17 | 0.02 |
| Gamma | 0.17 | 0.02 |
| Beta | 0.16 | 0.05 |
| Theta | 0.16 | 0.05 |

**Table S4.** Loadings and permutation-based p-values for brain variables contributing significantly to the second latent variable (LV2) identified by the PLS analysis. Brain loadings reflect the strength and direction of their contribution to the latent brain–behaviour association. Statistical significance was assessed using permutation testing (10,000 permutations), and only variables with p < .05 are reported.

| **Group** | **Sex**  **Female Male** | | **Mean ± SD**  **(age in months)** | **Mean ± SD**  **(age in years)** | **Range**  **(age in months)** |
| --- | --- | --- | --- | --- | --- |
| Cluster 1 | 17 | 43 | 138.42 ± 26.90 | 11.13 ± 2.26 | 97-204 |
| Cluster 2 | 6 | 18 | 171.83 ± 17.15 | 12.73 ± 1.42 | 148-201 |

**Table S5.**  Reports of the demographic characteristics of the sample across clusters, including age mean and standard deviation (in months and years), sex distribution and age range.

|  | **Variables** | **Mean ± SD Cluster 1** | **Mean ± SD Cluster 2** | **t** | **p-value  (FDR corrected)** |  |
| --- | --- | --- | --- | --- | --- | --- |
|  | Age (in months) | 138.42 ± 26.90 | 171.83 ± 17.15 | -6.78 | <0.001 | * |
| Regional Excitability | Bankssts L | 0.15 ± 0.04 | 0.08 ± 0.02 | 9.82 | <0.001 | * |
|  | Caudal middle frontal L | 0.13 ± 0.03 | 0.18 ± 0.03 | -5.92 | <0.001 | * |
|  | Caudal middle frontal l R | 0.10 ± 0.03 | 0.17 ± 0.04 | -8.31 | <0.001 | * |
|  | Cuneus R | 0.50 ± 0.08 | 0.37 ± 0.05 | 8.27 | <0.001 | * |
|  | Entorhinal L | 0.38 ± 0.06 | 0.27 ± 0.04 | 10.98 | <0.001 | * |
|  | Insula L | 0.36 ± 0.08 | 0.21 ± 0.07 | 8.44 | <0.001 | * |
|  | Insula R | 0.40 ± 0.08 | 0.22 ± 0.10 | 7.73 | <0.001 | * |
|  | Lateral orbito frontal L | 0.31 ± 0.04 | 0.39 ± 0.03 | -10.62 | <0.001 | * |
|  | Lateral orbito frontal R | 0.33 ± 0.06 | 0.40 ± 0.03 | -8.05 | <0.001 | * |
|  | Lingual L | 0.14 ± 0.03 | 0.17 ± 0.04 | -2.85 | 0.02 | * |
|  | Medial orbito frontal R | 0.31 ± 0.06 | 0.22 ± 0.05 | 6.55 | <0.001 | * |
|  | Pars orbitalis L | 0.25 ± 0.04 | 0.20 ± 0.02 | 7.32 | <0.001 | * |
|  | Pars orbitalis R | 0.19 ± 0.07 | 0.22 ± 0.11 | -1.20 | 0.43 |  |
|  | Postcentral R | 0.07 ± 0.02 | 0.09 ± 0.01 | -5.14 | <0.001 | * |
|  | Posterior cingulate L | 0.41 ± 0.05 | 0.35 ± 0.05 | 4.77 | <0.001 | * |
|  | Rostral anterior cingulate L | 0.51 ± 0.08 | 0.46 ± 0.06 | 3.20 | <0.01 | * |
|  | Supramarginal L | 0.07 ± 0.02 | 0.08 ± 0.02 | -2.07 | 0.10 |  |
| Global Fluidity | Theta | 0.03 ± 0.04 | 0.07 ± 0.05 | -4.00 | <0.001 | * |
|  | Alpha | 0.03 ± 0.04 | 0.08 ± 0.05 | -4.06 | <0.001 | * |
|  | Beta | 0.03 ± 0.04 | 0.08 ± 0.05 | -4.69 | <0.001 | * |
|  | Gamma | 0.03 ± 0.04 | 0.08 ± 0.05 | -3.76 | <0.01 | * |
| Self-report Questionnaires | Panas Positive | 16.28 ± 4.50 | 11.67 ± 3.80 | 4.77 | <0.001 | * |
| Neuro- psychological measures | Inhibition time NEPSY | 65.29 ± 13.37 | 51.67 ± 11.23 | 4.75 | <0.001 | * |
|  | Phonemic fluency BVN | 29.00 ± 8.43 | 35.25 ± 11.18 | -2.47 | 0.04 | * |

**Table S6.** Cluster differences in neural and behavioural variables (FDR-corrected p-values). Independent-samples t-tests were performed across clusters for neural and behavioural variables whose weights were significant in the PLS analysis. Measures were grouped into macro-domains (excitability, fluidity, questionnaires, neuropsychological measures). The table reports t-values, mean ± SD for each group, and FDR-corrected p-values.
